# Did a plant-herbivore arms race drive chemical diversity in *Euphorbia?*

**DOI:** 10.1101/323014

**Authors:** M. Ernst, L.-F. Nothias, J. J. J. van der Hooft, R. R. Silva, C. H. Saslis-Lagoudakis, O. M. Grace, K. Martinez-Swatson, G. Hassemer, L. A. Funez, H. T. Simonsen, M. H. Medema, D. Staerk, N. Nilsson, P. Lovato, P. C. Dorrestein, N. Rønsted

## Abstract

The genus *Euphorbia* is among the most diverse and species-rich plant genera on Earth, exhibiting a near-cosmopolitan distribution and extraordinary chemical diversity, especially across highly toxic macro-and polycyclic diterpenoids. However, very little is known about drivers and evolutionary origins of chemical diversity within *Euphorbia*. Here, we investigate 43 *Euphorbia* species to understand how geographic separation over evolutionary time has impacted chemical differentiation. We show that the structurally highly diverse *Euphorbia* diterpenoids are significantly reduced in species native to the Americas, compared to the Eurasian and African continents, where the genus originated. The localization of these compounds to young stems and roots suggest ecological relevance in herbivory defense and immunomodulatory defense mechanisms match diterpenoid levels, indicating chemo-evolutionary adaptation to reduced herbivory pressure.

**One Sentence Summary:** Global chemo-evolutionary adaptation of *Euphorbia* affected immunomodulatory defense mechanisms.

## Main Text

*Euphorbia* is among the most diverse and species-rich plant genera on Earth, exhibiting a near-cosmopolitan distribution and extraordinary chemical diversity among 2,000 species (*1–3*). The genus originated in Africa approximately 48 million years ago and through two single long-distance dispersal events 30 and 25 million years ago, expanded to the American continents (Fig. 1) (*1–2*). *Euphorbia* chemical diversity is characterized by an extraordinary diversity of macro-and polycyclic diterpenoids, biosynthetically derived from a head-to-tail cyclization of the tetraprenyl pyrophosphate precursor (*3, 4*). These compounds play an important ecological role as feeding deterrents and have shown exclusive occurrence and chemotaxonomic relevance in the plant families Euphorbiaceae and Thymelaceae (*3, 5–8*). However, the chemo-evolutionary transitions driving chemical diversity within *Euphorbia* are unknown. Here, we investigate the role of biogeography in the evolution of specialized metabolite diversity in 43 *Euphorbia* species, representing the genus’ global genetic diversity and biogeographic history across all continents.

**Fig. 1.**
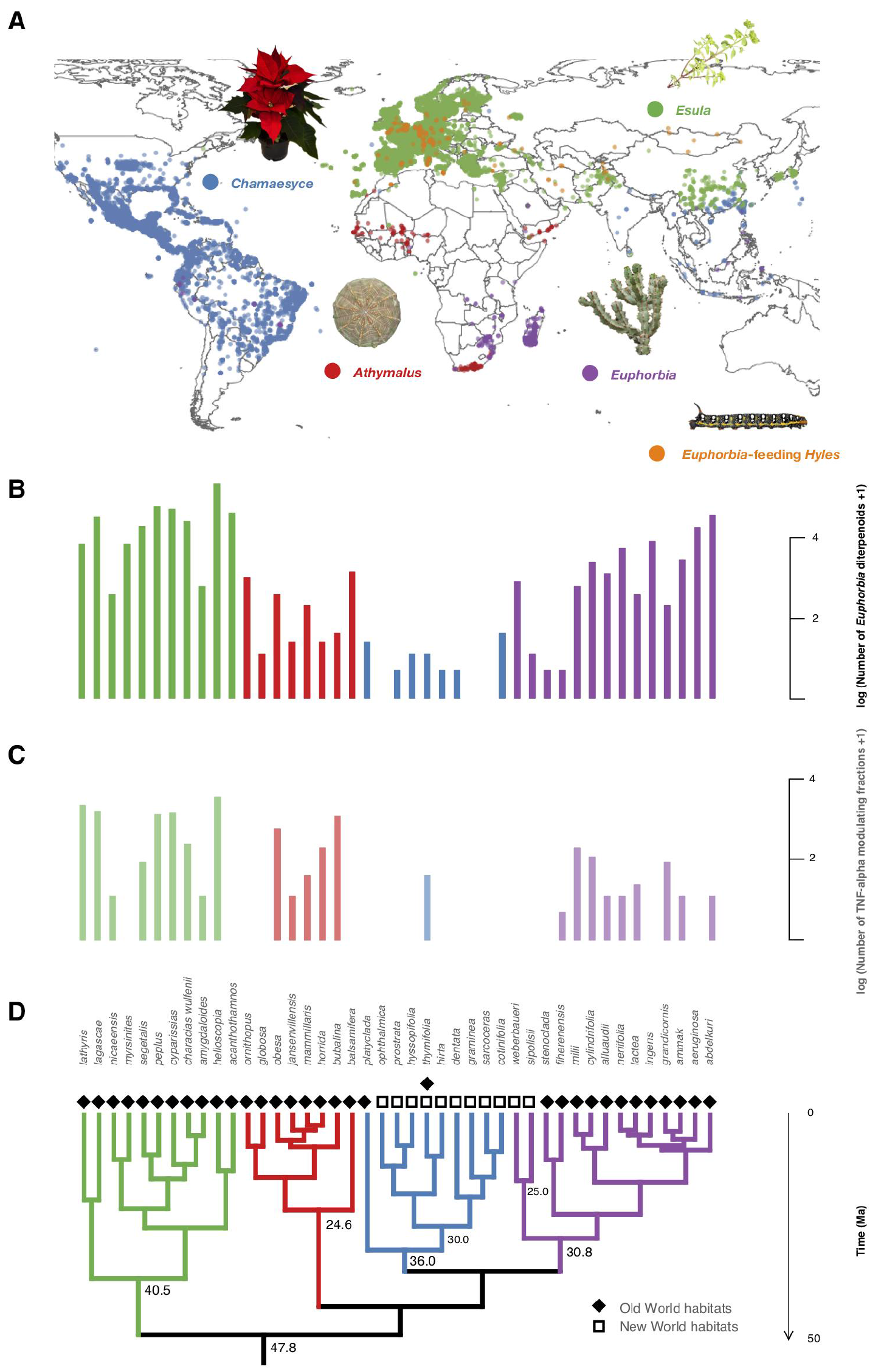
Biogeography, phylogenetic relationships, diterpenoid production and biological activities of representative *Euphorbia* species. **A**. Occurrences of *Euphorbia* species investigated chemically and *Euphorbia*-feeding *Hyles* moth larvae retrieved from GBIF and manually restricted to native areas **B**. Number of putatively annotated *Euphorbia* diterpenoids per species analyzed. **C**. Number of TNF-*α* modulating fractions per species analyzed. **D.** *Euphorbia* phylogenetic tree (50% majority rule consensus tree from Bayesian analysis of 11587 bps of DNA markers spanning all three plant genomes: chloroplast, mitochondrial, nuclear). Species of subgenus *Esula* exhibit a high number of biologically active diterpenoids and co-occur with larvae of *Euphorbia*-feeding *Hyles*, whereas the American radiation of subgenus *Chamaesyce* shows reduced *Euphorbia* diterpenoid production and TNF-*α* modulating activity. Subgeneric clades are highlighted with different colors: *Athymalus* (red), *Chamaesyce* (blue), *Esula* (green), *Euphorbia* (purple).

Coevolutionary theory suggests that an arms race between plants and herbivores yields increased specialized metabolite diversity (*9–11*). The evolution of a chemically different and biologically more active molecule increases a plant’s fitness by reducing the fitness of its predator (*10, 12*), and the probability of producing one or more biologically active compounds may increase with phytochemical diversity (*13*). To assess specialized metabolite diversity in relation to the evolutionary and biogeographic history of *Euphorbia*, we subjected extracts of 43 *Euphorbia* species to liquid chromatography tandem mass spectrometry (LC-MS/MS), created mass spectral molecular networks through Global Natural Products Social Molecular Networking (GNPS) (*14, 15*) and calculated the chemical structural and compositional similarity (CSCS) for all *Euphorbia* subgeneric clades (*16*). Our data show significantly higher chemical similarity among species of subgenus *Chamaesyce* compared to the mean chemical similarity among species of the remaining subgeneric clades (Fig. 2). The only species clustered in the chemogram (sharing high chemical similarity) are 8 out of 9 species representing the American clade within subgenus *Chamaesyce* (Fig. 2D). Consistent with the coevolutionary theory, the reduction of chemical structural diversity in these species suggests an adaptation to reduced herbivory pressure in the Americas during the biogeographic history of the genus.

**Fig. 2.**
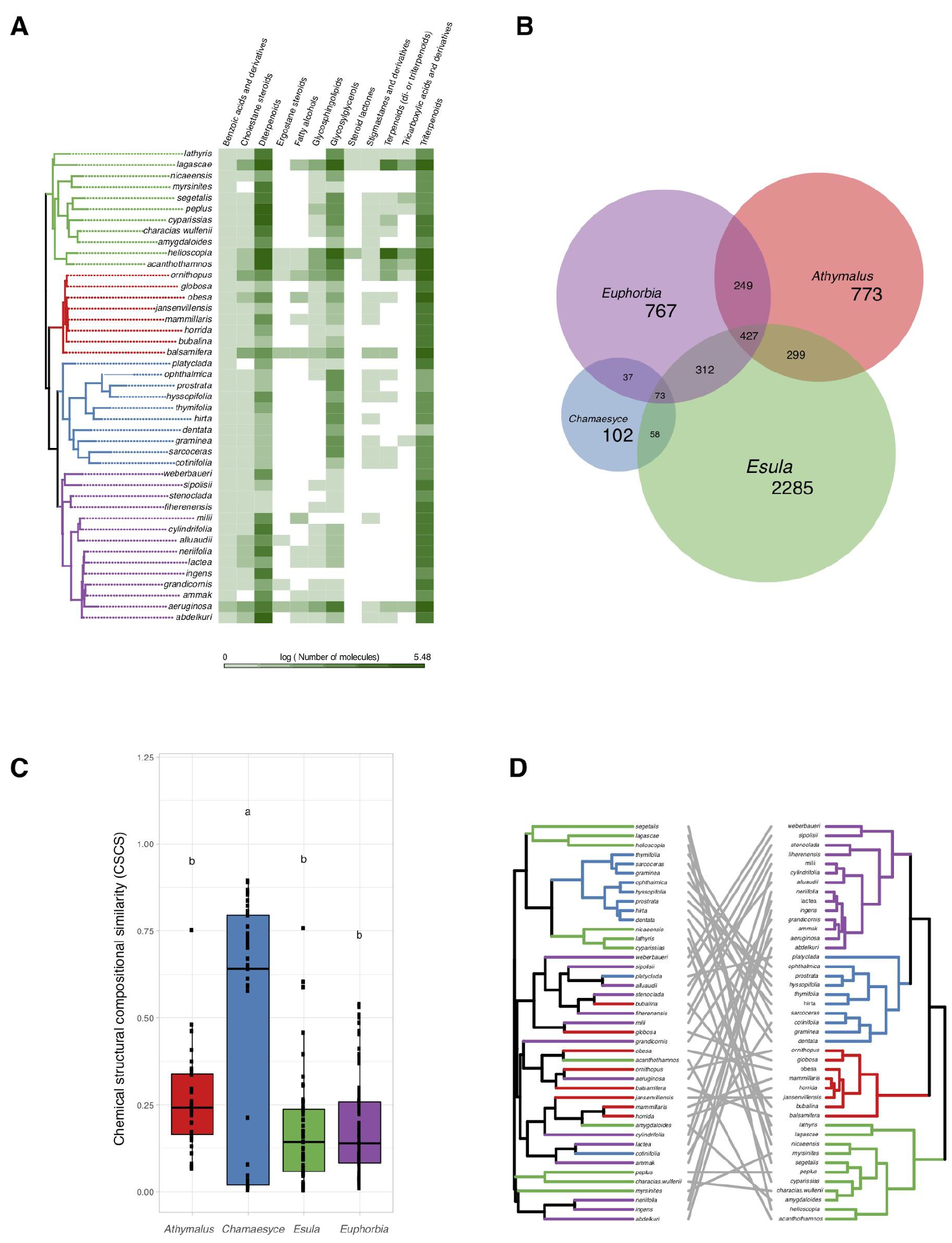
Specialized metabolite diversity in *Euphorbia*. **A**. Distribution of specialized metabolite classes on the *Euphorbia* phylogenetic tree (50% majority rule consensus tree from Bayesian analysis of 11587 bps of DNA markers spanning all three plant genomes: chloroplast, mitochondrial, nuclear). Chemical classes of *Euphorbia* specialized metabolites were identified using a mass spectrometry based workflow combining mass spectral molecular networking, *in silico* annotation, automated chemical classification and substructure recognition. **B.** Molecular features representing individual mass spectral molecular network nodes shared across species of *Euphorbia* subgeneric clades. **C**. Chemical similarity among *Euphorbia* subgeneric clades assessed using the chemical structural compositional similarity. Compared to subgenera *Athymalus*, *Esula* and *Euphorbia*, subgenus *Chamaesyce* exhibits very few chemically distinct features and high chemical structural compositional similarity. **D.** *Euphorbia* chemogram (left) and phylogenetic tree (right). The chemogram was generated using hierarchical cluster analysis on the pair-wise chemical structural and compositional dissimilarities of the tandem mass spectrometry data of the crude extracts using the complete agglomeration method. Phylogeny and chemogram show low overlap, suggesting that closely related *Euphorbia* species differ considerably in their chemistry.

To further understand the chemo-evolutionary relationships of *Euphorbia* at a molecular level we putatively identified major specialized metabolite classes by combining mass spectral molecular networking with *in silico* annotation tools (*17–19*), substructure recognition (*20, 21*) and automated chemical classification through ClassyFire (*22*). This resulted in annotated compound classes for over 30% of the compounds detected (*23, 24*) (Fig. S1 and S2, supplementary text). Our approach revealed many known *Euphorbia* diterpenoids as well as other metabolite classes. Out of six major structural classes of *Euphorbia* specialized metabolites (sesquiterpenoids, diterpenoids, cerebrosides, phenolics, flavonoids, and triterpenoids including steroids), four are found in our molecular networks (i.e., diterpenoids; triterpenoids including cholestane and ergostane steroids; steroid lactones and stigmastanes; and glycosylglycerols corresponding to cerebrosides). Additionally, *in silico* structure annotation suggests the presence of tricarboxylic and benzoic acids and derivatives as well as fatty alcohols and glycosphingolipids (Fig. 2A). Among the *Euphorbia* diterpenoids, we observe different skeletal types within the same molecular families (two or more connected components of a graph) (Fig. 3). Many *Euphorbia* diterpenoid backbone skeletons are isomeric, and their respective fragmentation spectra are highly similar (*25*). Nonetheless, we are able to distinguish different diterpene spectral fingerprints within a molecular family by mapping Mass2Motifs on the mass spectral molecular networks. Mass2Motifs correspond to common patterns of mass fragments and neutral losses, which are extracted using unsupervised substructure discovery of the MS/MS data through MS2LDA (*20, 21*). Combining *in silico* structure annotation, automated chemical classification, and MS2LDA, allows us to putatively identify chemical classes within the mass spectral molecular networks, as well as chemical subclasses within molecular families (Fig. 3, Fig. S2). Consistent with previous observations of *Euphorbia* diterpenoids exhibiting anti-herbivore biological activity (*5–8*) and the low chemical diversity exhibited by subgenus *Chamaesyce*, we find very few to no *Euphorbia* diterpenoids in representatives of the American radiation of subgenus *Chamaesyce* (Fig. 1). Although subgenus *Chamaesyce* includes the largest American radiation within the genus *Euphorbia*, there is also a smaller American radiation within subgenus *Euphorbia* (Fig. 1). The two investigated representatives of this clade contain intermediate amounts of *Euphorbia* diterpenoids. The clade is estimated to have originated approximately 5 million years later than the American radiation within subgenus *Chamaesyce* (*1*), which could suggest less time for adaptation to reduced herbivory pressure.

**Fig. 3.**
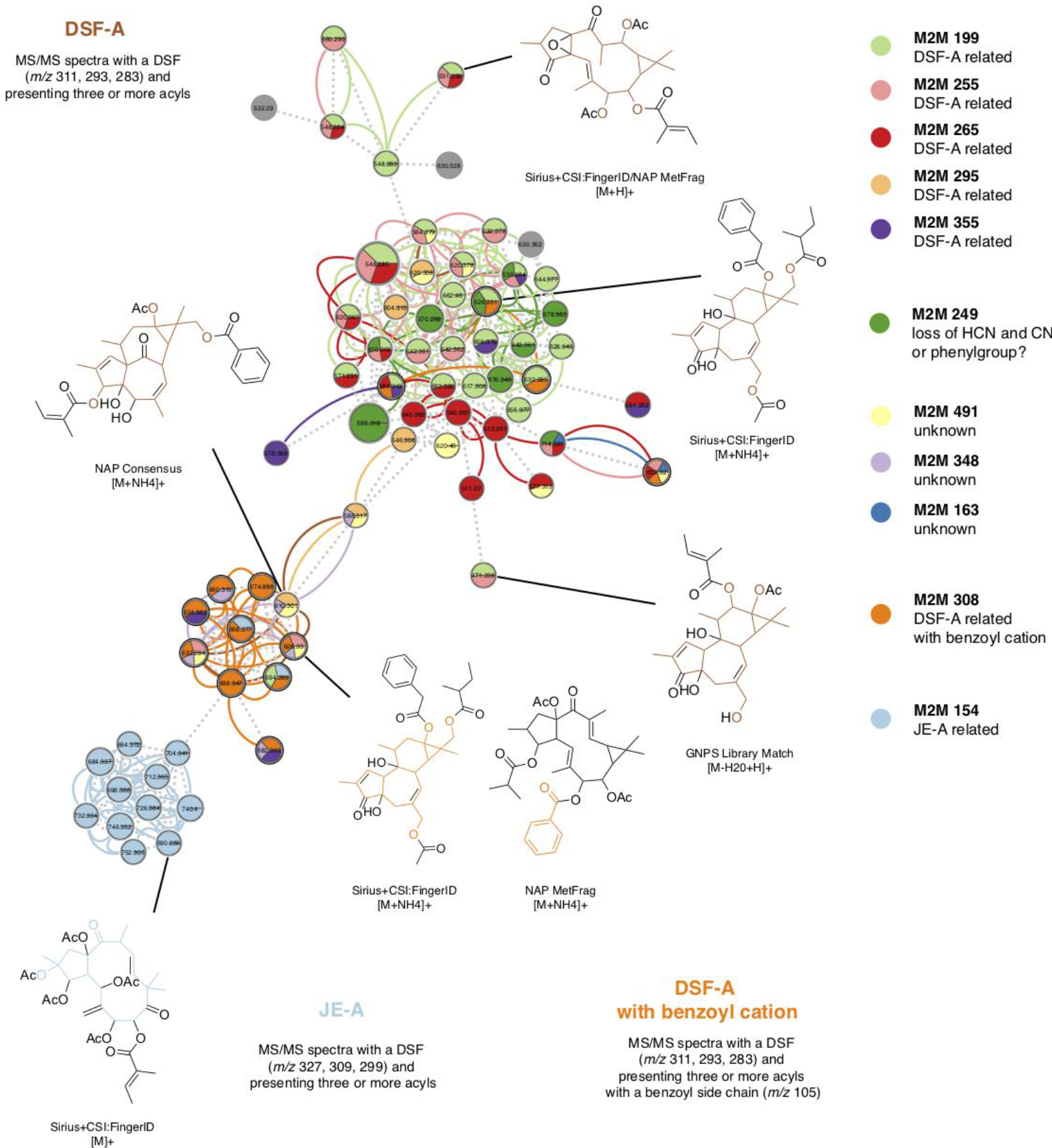
Putative identification of chemical compound classes. We putatively identified compound classes within the mass spectral molecular networks by combining *in silico* annotation with automated chemical classification and substructure recognition (MS2LDA). *Euphorbia* diterpenoids exhibit many isoforms, therefore different diterpene backbone skeletons were found within the same molecular family. Matching substructures (Mass2Motifs) associated with diterpenoid substructures obtained from matches to reference spectra and *in silico* structure annotation enabled the identification of different diterpene spectral fingerprints clustered within one molecular family. Node size represents the total ion current (TIC) of all samples analyzed, edge colors represent different substructures (Mass2Motifs) that are shared across different nodes and dotted lines connecting the nodes represent the cosine score. M2M: Mass2Motif, DSF-A: Diterpene spectral fingerprint type A, JE-A: Jatrophane ester type A.

To understand where the *Euphorbia* diterpenoids are produced within the plants, we dissected four species representing the four subgeneric clades into approximately 20 sections(*23*) (Fig. S10, Fig. S11). Mass spectrometric investigation revealed that diterpenoids are primarily found in the roots, in representatives of subgenera *Euphorbia* and *Athymalus* (*E. milii* var. *hislopii* and *E. horrida,* Fig. 4, Fig. S3, Fig. S6-S9). In the European subgenus *Esula* (*E. lathyris*), diterpenoid production is also pronounced in other plant parts, such as the young stems (Fig. 4, Fig. S5). Consistent with the lower chemical diversity reported above, diterpenoid production is reduced or absent from most sections throughout the whole plant in *E. hirta*, a representative of the American clade within subgenus *Chamaesyce* (Fig. 4, Fig. S4, Fig. S10). Compartmentalization of the diterpenoids to mainly young stems and roots underpins their function as anti-feeding molecules (*5–8*).

**Fig. 4.**
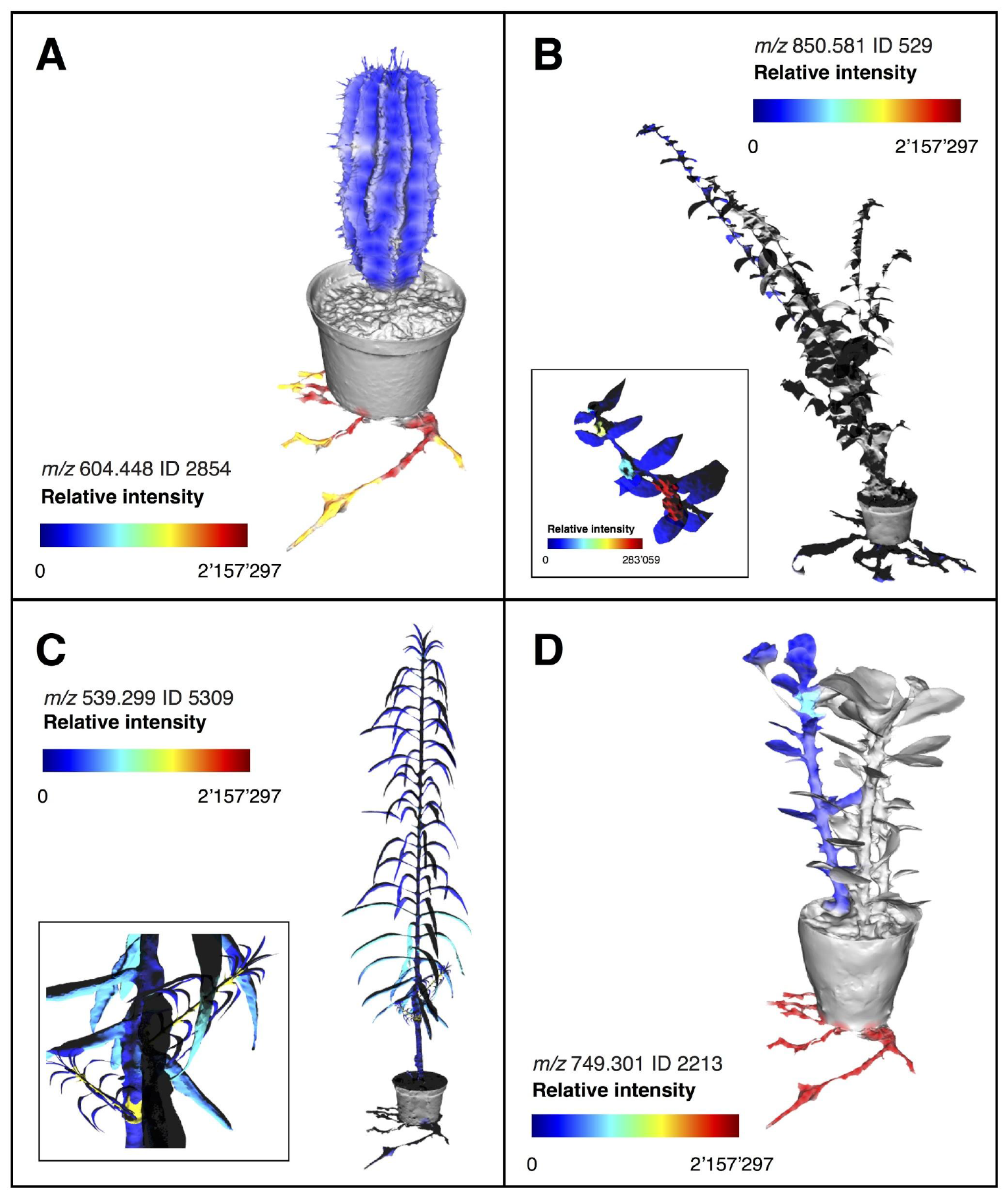
Molecular maps of selected Euphorbia diterpenoids across representatives of each subgeneric clade. Relative intensity of LC-MS molecular features annotated as *Euphorbia* diterpenoids through spectral matching **A**. *Euphorbia horrida*, subgenus *Athymalus*, **B**. *Euphorbia hirta*, subgenus *Chamaesyce*, **C**. *Euphorbia lathyris*, subgenus *Esula* and **D**. *Euphorbia milii* var. *hislopii*, subgenus *Euphorbia*. For interactive cartographical snapshots see URL S1, links 1–15. The 3D images are for illustrative purposes only and do not represent exact locations of sample collection.

As anti-herbivore activity cannot be directly tested from the past continental transition, we set out to test a bioactivity that could reflect a chemical defense mechanism. One strategy of defense invoked by plants to overcome their sessile habit, is through immunomodulatory effects on herbivores. *Euphorbia* diterpenoids are known to exhibit immunomodulatory activities through the selective modulation of protein kinase C (PKC) (*26, 27*). Therefore, we evaluated the modulation of PKC by measuring the capacity of small extract fractions, corresponding to compounds or compound groups in each of the 43 *Euphorbia* species, that modulate *in vitro* TNF-α release from peripheral blood mononuclear cells (PBMCs). To pinpoint the role of *Euphorbia* diterpenoids involved with TNF-α modulating properties, we tested for correlation between the number of bioactive fractions and the number of molecules within the previously annotated chemical classes using phylogenetic generalized least squares regression analysis (PGLS). The association between the number of TNF-α modulating fractions and the number of *Euphorbia* diterpenoids is significant (*P*-value: 0.02) (Fig. 1B, Fig. 1C, Table S1). Besides the *Euphorbia* diterpenoids, we also observe significant associations (*P*-value<0.05) between the number of TNF-α modulating fractions and the number of overall diterpenoids and glycosyl glycerols with the best fit observed for the *Euphorbia* diterpenoids (Fig. 1B, Fig. 1C, Fig. S12, Table S1), supporting our hypothesis of the ecological function of *Euphorbia* diterpenoids as immunomodulatory defense molecules.

To evaluate the possibility of *Euphorbia* diterpenoids being produced as a response to local plant-predator interactions, we compiled a dataset of known *Euphorbia* herbivores from the literature (*23*). Several hawkmoth species of the genus *Hyles* were found to be highly specialized predators of *Euphorbia* (*5, 28*). They have been shown to exhibit host specificity and to tolerate the highly toxic *Euphorbia* diterpenoids, which they reuse as a defense strategy against their own predators by regurgitating plant material from the gut (*5, 8*). Native species distribution data suggests a close co-occurrence of *Euphorbia*-feeding *Hyles* species with the chemically highly diverse and biologically active European species of subgenus *Esula* (Fig. 1A), and an absence in the American habitats of the chemically less diverse and biologically little active representatives of subgenus *Chamaesyce*. However, our data also suggests that African members of subgenus *Athymalus* and subgenus *Euphorbia* occurring in Southern Africa and Madagascar, outside of the distribution range of *Euphorbia*-feeding *Hyles*, produce a high diversity of feeding deterrent diterpenoids (Fig. 1). Thus, we speculate, that previously not described (or extinct) generalist or specialist *Euphorbia*-feeding herbivores occur (or occurred) in these regions, which contributed to maintaining adaptive pressure. The black rhinoceros, *Diceros bicornis* L. distributed in the southern African subregion, for example, was found to feed often and extensively on African *Euphorbia* species of subgenus *Euphorbia* (*29*). Anecdotal evidence of the lack of specialized herbivores in the Americas supports our results. A single hawkmoth species, *Hyles euphorbiae*, was only introduced recently to North America (*5*), where it was used as a host-specific enemy and biological control for the reportedly highly invasive European species of subgenus *Esula* (*E. cyparissias* and *E. esula*), lacking predators in the newly occupied habitats. Furthermore, although the poinsettia (*E. pulcherrima* Willd. ex Klotzsch), a very well-known house plant and American representative of subgenus *Chamaesyce,* is notorious for its “extreme toxicity” among the general public, toxicity remains unconfirmed in the clinic, as 92.4% of patients exposed to the plant did not develop adverse effects (*30*), corroborating the low chemical diversity and immunomodulatory activity observed here.

In contempt of the limited knowledge about *Euphorbia* herbivory, the remarkable immunomodulatory activities of the *Euphorbia* diterpenoids provide an indirect way to assess chemical defense against predators. The differential biosynthesis of diterpenoids in species geographically separated through evolutionary time, suggests differential exposure to herbivory during the biogeographic history of the genus. Indeed, there are no known herbivores of the American species, while specialized herbivores are well documented for the European and African species (*8, 29*). The mechanism of predator tolerance is not known, but the presence of specialized herbivores is consistent with our results and hypothesis that the greater the diversity of herbivores feeding on a plant species, the more immunomodulating molecules the plants produce.

## Acknowledgments

The authors thank Ivan Protsyuk and Theodore Alexandrov (EMBL) for their support in using the KNIME workflow Optimus and MassFuser, Arife Önder for technical assistance with fractionation, Martin Aarseth-Hansen and Gry Bastholm for assistance with collecting plant material and co-workers of LEO Pharma for technical assistance with biological assays. Kevin Nguyen, Brandon Ross and Kirandeep Chhokar (UCSD) are thanked for assistance with preparing plant extracts for the 3D mass spectral molecular cartography;

## Author contributions

ME, LFN, CHSL, OMG, HTS, NN, DS, PCD and NR designed the study. ME and KM collected and sampled *Euphorbia* species from the Botanical Garden at the Natural History Museum of Denmark. GH and LAF collected seeds of Brazilian *Euphorbia* species and performed taxonomic circumscription and identification. ME prepared the plant extracts. LFN performed LC-MS/MS analysis of the pooled *Euphorbia* extracts. ME performed LC-MS/MS analysis for the 3D mass spectral molecular cartography. ME and LFN performed mass spectral molecular networking analysis. JJJVDH performed MS2LDA analysis. LFN, JJJVDH and ME annotated Mass2Motifs on MS2LDA. ME and JJJVDH developed and performed the (semi-)automated annotation workflow by combining mass spectral molecular networking with MS2LA, *in silico* annotation and ClassyFire. RRS developed dereplication strategies through NAP and provided support for data analysis. NN, PL, DS and ME designed the high-resolution bioactivity study. ME performed the high-resolution bioactivity study and analysed data together with DS, NN and PL. ME generated the phylogenetic hypothesis, performed comparative analyses and wrote all scripts used for data analysis. ME wrote the manuscript together with LFN, NR and PCD. All authors discussed the results and commented on the manuscript.

## Data and materials availability

All data, code, and materials to understand and assess the conclusions of this research are available in the main text, supplementary materials, and via https://github.com/DorresteinLaboratory/supplementary-GlobalEuphorbiaStudy. LC-MS/MS data are publicly accessible on GNPS under the MassIVE accession no. MSV000081082 and MSV000081083.

## Supplementary Materials

Materials and Methods

Supplementary Text

Table S1

Fig S1-S12

URL S1

Data S1-S2

References (*31–51*)

